# Contrastive learning of protein representations with graph neural networks for structural and functional annotations

**DOI:** 10.1101/2022.11.29.518451

**Authors:** Jiaqi Luo, Yunan Luo

**Author notes:** Preprint of an article published in Pacific Symposium on Biocomputing © 2022 World Scientific Publishing Co., Singapore, http://psb.stanford.edu/. © 2022 The Authors. Open Access chapter published by World Scientific Publishing Company and distributed under the terms of the Creative Commons Attribution Non-Commercial (CC BY-NC) 4.0 License.

## Abstract

Although protein sequence data is growing at an ever-increasing rate, the protein universe is still sparsely annotated with functional and structural annotations. Computational approaches have become efficient solutions to infer annotations for unlabeled proteins by transferring knowledge from proteins with experimental annotations. Despite the increasing availability of protein structure data and the high coverage of high-quality predicted structures, e.g., by AlphaFold, many existing computational tools still only rely on sequence data to predict structural or functional annotations, including alignment algorithms such as BLAST and several sequence-based deep learning models. Here, we develop PenLight, a general deep learning framework for protein structural and functional annotations. PenLight uses a graph neural network (GNN) to integrate 3D protein structure data and protein language model representations. In addition, PenLight applies a contrastive learning strategy to train the GNN for learning protein representations that reflect similarities beyond sequence identity, such as semantic similarities in the function or structure space. We benchmarked PenLight on a structural classification task and a functional annotation task, where PenLight achieved higher prediction accuracy and coverage than state-of-the-art methods.

## 1. Introduction

With the decrease in the cost of sequencing technology, protein sequence data have been accumulated to an ever-increasing amount. How to characterize those amino acid sequences with structural and functional annotations is a long-standing and challenging problem in bioinformatics. The community has long been interested in developing computational tools to infer protein functions from their sequences, ranging from BLAST,^1^ profile hidden Markov models (pHMM),^2^ and several other popular methods.^3–6^ Despite the success of these tools in inferring protein functional annotations, such as the Gene Ontology (GO) terms and Enzyme Commission (EC) numbers, the whole protein universe is still sparsely annotated. For example, in Pfam, a popular protein family database, it was reported that one-third of bacterial proteins cannot be annotated by alignment approaches.^7^

Recently, deep learning (DL) has emerged as a promising approach to complement traditional tools to expand protein annotations and has gained impressive success. For instance, Bileschi et al. develop a deep neural network to predict protein functional labels, which was adopted by the Pfam database to expand its coverage by > 9.5%.^8^ Other successful applications of DL in protein annotation include structure fold recognition,^9^ GO term prediction^10^ and EC number predictions.^11^ Another notable trend along this line is protein language models (PLMs), which learn rich representations that encode intrinsic biophysical, evolutionary, and structural properties of proteins from large-scale unlabeled protein sequence data. PLMs have been found to substantially improve prediction accuracy for many protein structure and function prediction problems.^12^

It is believed that protein sequence determines protein structure, which dictates function. Knowing the three-dimensional (3D) information of protein structures can be useful for protein function prediction because structures are more conserved than sequences and more directly related to functions such as protein binding. However, due to the limited availability of solved protein structure data, most existing methods for functional annotations are trying to directly predict functions from sequences, assuming that proteins sharing high sequence similarity will have the same set of functions. This assumption may not always hold, as it has been found that proteins with similar structures can have seemly random sequence similarity. Fortunately, with advances in biotechnology such as cryo-EM,^13^ the number of solved protein structures is constantly increasing.^14^ The structure coverage is further improved by the high-quality structures predicted by DL models such as AlphaFold.^15^ Remarkably, in August 2022, DeepMind released 200M AlphaFold’s predicted structures, covering nearly every known protein on the planet. In parallel, the machine learning community has made great advancements in developing graph neural networks (GNNs) for modeling graph data, which have resulted in successful applications such as AlphaFold.^15^ Despite the new opportunity offered by the largely available solved and predicted structures and the advancements in GNNs, integrating structure data and graph DL has not been widely exploited for protein functional and structural annotations.

The supervised learning paradigm has been a popular choice in previous deep learning methods for predicting protein functions, in which the protein sequence is directly mapped to the class output. This paradigm faces the challenge of class imbalance. For example, many Pfam families contain relatively few sequences, which makes it difficult for supervised models to predict because the training objective is dominated by the major Pfam classes. Another paradigm called contrastive learning has recently gained interest in the machine learning community.^16^ Instead of directly mapping sequences to functions, contrastive learning optimizes a latent embedding space where sequences with similar functions are pulled together, while sequences of different functions are pushed away. The ProtTucker model developed by Heinzinger et al.^17^ was among the first attempts of using contrastive learning for protein annotation, but the model only predicts protein structural annotations from protein sequence information. Extending contrastive learning to integrate structure data has not been explored for protein structural and functional annotations.

Here, we present PenLight (<u>P</u>rot<u>e</u>in co<u>n</u>trastive <u>l</u>earn<u>i</u>ng with <u>g</u>rap<u>h</u> neural network for anno<u>t</u>ation), a contrastive deep learning model for protein structural and functional annotations. PenLight models protein 3D structure as a graph and uses a GNN to learn structure-aware representations for the input protein. A major innovation of our work is using contrastive learning for refining the learned protein representations so that the semantic similarity of protein structures or functions can be reflected in the embedding space. We demonstrate PenLight’s applicability using a structure classification task (fold classification) and a functional annotation task (EC number prediction). On both tasks, PenLight outperformed existing methods, including alignment algorithms such as BLAST and previous deep learning approaches. We observed that PenLight was able to achieve high prediction accuracy as well as high coverage. We expect PenLight to be used as a general deep learning framework for protein annotations.

## 2. Materials and Methods

### Overview of PenLight

In this work, we develop PenLight, a graph neural network trained with contrastive learning, for predicting protein structural and functional annotations. As an overview (Fig. 1), PenLight receives the three-dimensional structure of a protein as input and represents it as a graph, where the graph’s nodes are protein residues, and the edges encode the spatial proximity of residues. Protein language model embeddings and a set of geometric features (e.g., distance and orientation) derived from the input structure are used to initialize the node and edge features. PenLight then employs a contrastive learning scheme to learn a vector representation for each protein, such that the representations of structurally/functional similar proteins are pulled together while dissimilar proteins are pushed apart. PenLight then transfers the known annotations of a protein to an unlabeled protein if their representation distance is below a threshold. The source code of PenLight is available at https://github.com/luo-group/PenLight.

**Fig. 1.**
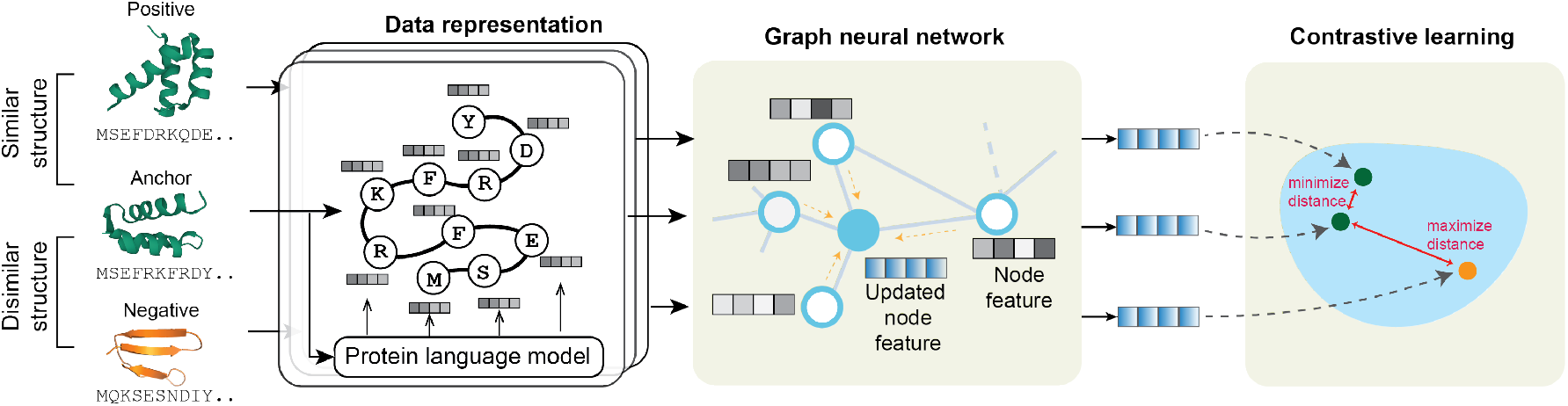
Schematic illustration of PenLight.

#### 2.1. Tasks and Datasets

We showcase the applicability of PenLight using a structure classification task and a functional annotation task. Specifically, we train separate PenLight models to predict the structure classification code in the CATH database and the enzyme class (EC number) of a protein. Both CATH codes and EC numbers are four-level classification systems that characterize different levels of similarities of proteins, as described below.

##### Structure classification

We utilize the CATH dataset,^18^ an expert-curated database that classifies 3D protein structures from the Protein Data Bank (PDB) database^14^ into a hierarchical classification system. We downloaded and processed the structures from CATH following Heinzinger et al..^17^ Each protein structure is assigned with a label (CATH code) at the Class (C), Architecture (A), Topology (T), and Homologous superfamily (H) levels, respectively. Intuitively, higher levels (H>T>A>C) contain proteins that are more similar in their 3D structure.

##### Functional annotation

We choose the Enzyme Commission number (EC number) prediction as an example of functional annotation tasks. Similar to CATH, EC number is also a four-level numerical classification scheme for enzymes, which assigns each enzyme with a label based on the chemical reactions it catalyzes. We downloaded structures annotated with EC numbers in the PDB database following a previous study.^19^ While there exist promiscuous enzymes that are labeled with more than one EC number, most enzymes are labeled with only a single EC number. Therefore, we only consider the top-1 predictions when evaluating different prediction methods in this work.

#### 2.2. Protein Structure Representations

The structure data of a protein contains the three-dimensional (3D) coordinates of atoms of the protein structure. Here, we focus on the *C_α_* atoms of the backbone and use them to represent the residues of a protein. We denote the coordinates of those *C_α_* atoms as 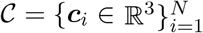, where *N* is the number of residues. We represent the structure as a graph 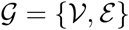, where the node set 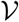 contains the residues and the edger set 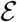 indicates the residue contacts, which is defined by a distance cutoff of 8Å between pairwise *C_α_* atoms.

To improve the expressiveness of the structure representation, we also associated features to each node and edge in the graph 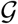. We built a series of features that are invariant to rotations and translations following a previous study.^20^ For the node feature ***v_i_*** of residue *i*, we used the per-residue embeddings generated by ESM-1b^12^ or ProtT5,^21^ protein language models (PLM) that are trained on millions of protein sequences using unsupervised representation learning. It has been shown that PLM can boost the prediction accuracy for protein function and structure predictions.^12,22^ We used ProtT5 embeddings for the structure classification task following Heinzinger et al.^17^ and ESM-1b for the functional annotation task, as we found in nested cross-validation that this resulted in a better performance. For the edge between residues *i* and *j*, we concatenated multiple features ***e**_ij_* = [(***c**_j_* – ***c**_i_*)/||***c**_j_* – ***c**_i_*||_2_; RBF(||***c**_j_* – ***c**_i_*||_2_); *E_pos_*(***c**_j_* – ***c**_i_*)], where the first term is the unit direction vector, the second term is the pairwise distance lifted into radial basis functions (RBFs), and the third term is the sinusoidal encoding of the relative distance and direction between the two residues.

#### 2.3. Graph Neural Network

Now we introduced the GNN architecture used in PenLight. We used a modified version of graph attention network (GAT)^23,24^ as our backbone model. Given the structure graph 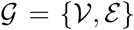 of the input protein, GAT applies *L* layers of graph convolution operations that transform 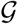 to an embedding 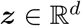. The *ℓ*-th layer transforms residue *i*’s embedding 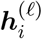 to an updated embedding 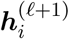 by aggregating the information from residue *i* and its neighbor residues: 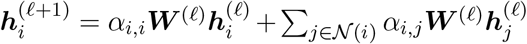, where 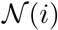 is the set of neighbor nodes of node *i*, **W** are learnable weights of the GNN, and *α_ij_*’s are attention weights used to adaptively aggregate embeddings from node *i*’s neighbors. The embedding 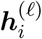 is initialized using the node feature ***v**_i_* for *ℓ* = 0. The attention weights are computed as (the superscript of layer index ℓ is omitted for simplicity):

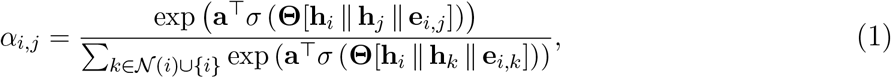

where ***a*** and **Θ** are learnable weights, ∥ is vector concatenation, and *σ*(·) is the Leaky ReLU activation function. PenLight used two stacked GAT layers with ReLU activation to transform the initial node features into 512-dimensional vectors 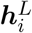 for each amino acid. A global mean pooling layer was used after the GNN to aggregate the embeddings of all amino acids into embeddings into a single embedding 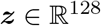, representing the input protein.

#### 2.4. Contrastive Learning

We applied contrastive learning to optimize the GNN model in PenLight, which directly optimizes an embedding space such that proteins with the same structural or functional category are located together in the embedding space. The GNN model receives a triplet of proteins (represented as graphs) as input each time, i.e., an anchor protein *x_a_*, a positive protein *x_p_* that is structurally/functionally similar to *x_a_*, and a negative protein *x_n_* that is structurally/functionally dissimilar to *x_a_*. The objective of contrastive learning is to learn an embedding function (parameterized by the GNN) 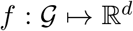 such that the distance between the positive pair is smaller than that of the negative pair: *d*(*f*(*x_a_*), *f*(*x_p_*)) < *d*(*f*(*x_a_*), *f*(*x_n_*)), where *d*(·, ·) is a distance function (e.g., Euclidean distance) defined on the embedding space.

##### Triplet sampling

How to sample the triplets is the key to learning a well-organized embedding space. Since both CATH codes and EC numbers are organized in hierarchical tree structures with four levels, and each label is represented as a four-digit number from coarse to fine (e.g., EC: 3.2.1.2), we adopted a hierarchical sampling strategy^17^ to randomly sample the triplets (*x_a_, x_p_, x_n_*) for both tasks. More specifically, during training, we sampled each protein in the training set as the anchor protein *x_a_*. For each anchor protein we first randomly chose a similarity level *γ* ∈ {1, 2, 3, 4}. Then a different protein with the same label up to the *γ*-th digit was sampled as the positive protein *x_p_*, and another protein with a different digit at the *γ*-th level but the same digit at the (*γ* – 1)-th level was sampled as the negative protein *x_n_*. For example, if we sampled an anchor protein with CATH label 2.20.25.20 and we randomly chose the similarity level *γ* = 2 (the Architecture level), the positive protein should be randomly sampled from proteins with CATH label of type 2.20.*.* (i.e., having the same first two digits) and the negative should be randomly sampled from those with CATH label 2.*a*.*.* where *a* is not 20 (share the same Class code but different Architecture code).

##### Hard negatives/positives mining

Previous studies^25^ have shown that another key to successful contrastive learning is the balance between the triviality and the hardness of the sampled triplets. Here, we further enhance the triplet samples by mining hard negatives and positives to improve the performance of contrastive learning, as did in Heinzinger et al..^17^ During training, we utilized the batch-hard^25^ technique inside each mini-batch. After getting a mini-batch of hierarchical sampled triplets, we shuffled all the anchor, positive and negative proteins in the mini-batch and applied hierarchical sampling in these proteins but with one more criterion that the positive had the maximum Euclidean embedding distance with the anchor among all the positive candidates selected under hierarchical sampling while the negative had the minimum distance with the anchor.

##### Training

During the model training, PenLight receives the sampled triplet as input and uses the GAT model to transform them into *d*-dimensional embeddings (the three GATs for anchor, positive and negative shared the same set of parameters). Based on inner-loop cross-validation results, the embedding size was set to 128 in the CATH classification task and 256 in the EC number prediction task. We used the soft margin loss as the objective to train PenLight: 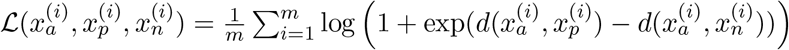, where *m* is the dimension of the output embeddings, *d*(·, ·) is the Euclidean distance between embeddings. Adam with an initial learning rate 1e-4 and a weight decay of 1e-4 was used as the optimizer. Early stopping was also applied to avoid overfitting. We set the batch size to 256.

#### 2.5. Inference and Evaluation

Since contrastive learning yielded a vector embedding instead of a direct label for each input protein, the final inference would be performed in a query-lookup manner. Given a lookup set 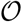, which contains proteins with known (structural or functional) labels, and a query (unlabeled) protein *q* that we would like to infer labels for, PenLight projects all proteins in 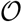 and *q* into the same embedding space. We call a protein 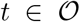 a “hit” for the query protein *q* if their Euclidean embedding distance is below some threshold *δ*. We can then infer the annotations for the query *q* by transferring the annotations of all hit proteins, i.e., 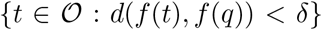, to the query *q*. The inference for individual query protein is very efficient since it only requires a single forward pass of the graph neural network and a distance comparison, both of which are matrix or vector operations that can be accelerated on GPUs. In practice, we found that the average inference time per protein was 0.68 seconds for CATH classification and 0.04 seconds for EC number prediction. We also observed that the prediction accuracy can be improved by an ensemble approach, i.e., two replicas of PenLight were trained on the same data, and the average distance given by them was used to find the hit proteins.

To evaluate the performance of PenLight and other baseline methods, we computed the accuracy, precision, recall, and F1 scores for each class (CATH code and EC number) and then average the metrics over all classes (i.e., macro-averaged metrics). Some baseline methods use a confidence threshold to decide whether to predict the annotations for a query protein (e.g, the E-value in BLAST). For those methods, we count it as a wrong prediction if the model does not predict any annotation for a query protein, unless otherwise specified.

## 3. Results

### 3.1. Performance on Structure Classification

We downloaded the CATH-S100 dataset (123k proteins, clustered based on identity 100%) from the CATH database (v4.3),^18^ including both the structure data and their CATH code labels. We followed the study by Heinzinger et al.^17^ to split the dataset into four splits, namely, the training set (~71k proteins), validation set (196 proteins), lookup set ~74k proteins), and test set (208 proteins). The median number of samples per CATH class is 2. The splits were created using the clusters generated by MMseqs2^6^ such that any sequence in the training set does not share > 20% sequence identity to any protein in the validation or test set. To directly test PenLight’s ability to transfer structural annotations from labeled proteins to unlabeled proteins, an independent lookup set that contains ~74k proteins was also created. Redundant sequences shared by the test set and lookup set were also removed. We compared PenLight with different types of baseline methods for structure classification, including sequence alignment algorithm (BLASTp^1^), unsupervised PLMs (ESM-1b^12^ and ProtT5^21^), and the state-of-the-art contrastive learning method for structural annotation (ProtTucker^17^). For PLM baselines, we predicted the annotations for test proteins by applying an unsupervised *k*-nearest neighbor classifier with *k* = 1 or a supervised multi-class classifier (ProtT5-sup) on PLM representations.

We observed that PenLight consistently outperformed other methods when evaluated using several metrics (Table 1). First, we noticed that the information-rich features used in PenLight are extremely useful for predicting the CATH code. For example, PenLight achieved substantial improvements (+120% in accuracy and +146% in F1) compared to BLASTp, which only uses the raw amino acid sequences to perform sequence comparison. Second, our results also suggested the benefits of contrastive learning (CL) in PenLight. The PLM embeddings, used as the initial features in PenLight, were trained purely on sequence data and may not explicitly capture structure properties. However, the contrastive learning used in PenLight is able to refine the PLM embeddings to be discriminative and structure-aware by utilizing the CATH hierarchy. This is demonstrated in the clear distribution separation of structurally similar and dissimilar proteins in the embedding space (Fig. 2a). The well organized embedding space also translated into performance improvement, where PenLight boosted PLM’s F1 score from 0.25+ (for ESM1-1b and ProtT5) to 0.37 (Table 1). These improvements suggested that contrastive learning is effective in learning representations that reflect the semantic similarities in the label space (e.g., the CATH classification here). Finally, we observed that PenLight also outperformed the state-of-the-art method ProtTucker that only considered sequence data as input, suggesting that incorporating the 3D structure information as input is useful for predicting the CATH classification of proteins. Overall, these results demonstrated PenLight’s improved prediction performance in predicting the structural annotations of proteins.

**Fig. 2.**
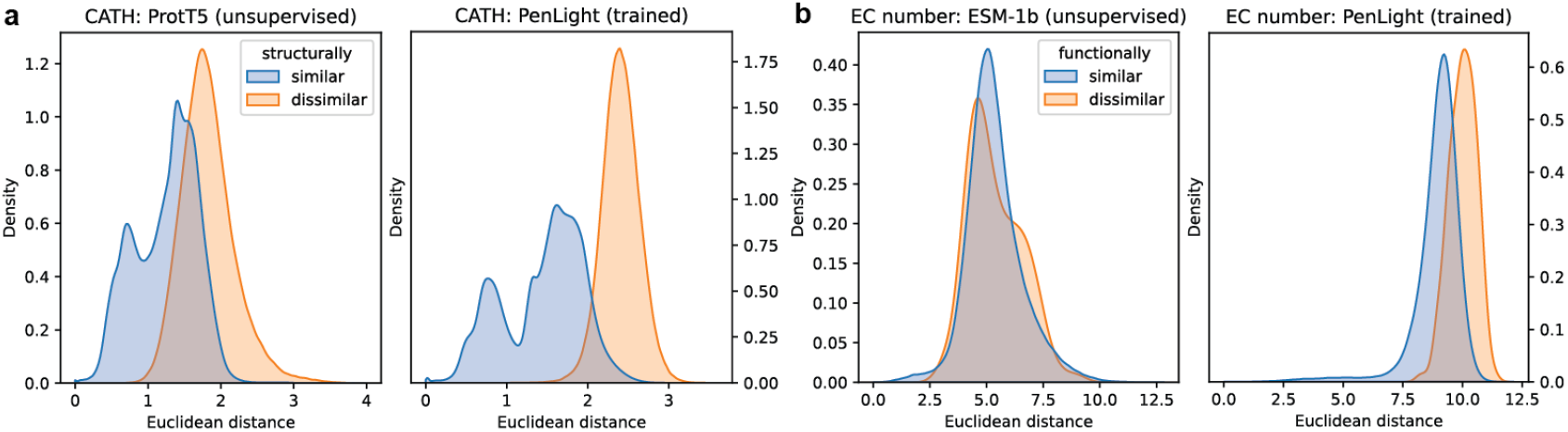
PenLight separated structural or functional similar proteins from dissimilar ones in the embedding space. We consider two proteins are *structurally* similar if they are assigned with the same third-level but different fourth-level CATH codes, and two proteins are *functionally* similar if they are assigned with the same second-level but different fourth-level EC numbers. Euclidean embedding distances learned by PenLight and two PLMs were visualized for similar and dissimilar proteins in the training sequences of (a) the CATH dataset and (b) the EC number dataset.

**Table 1.**
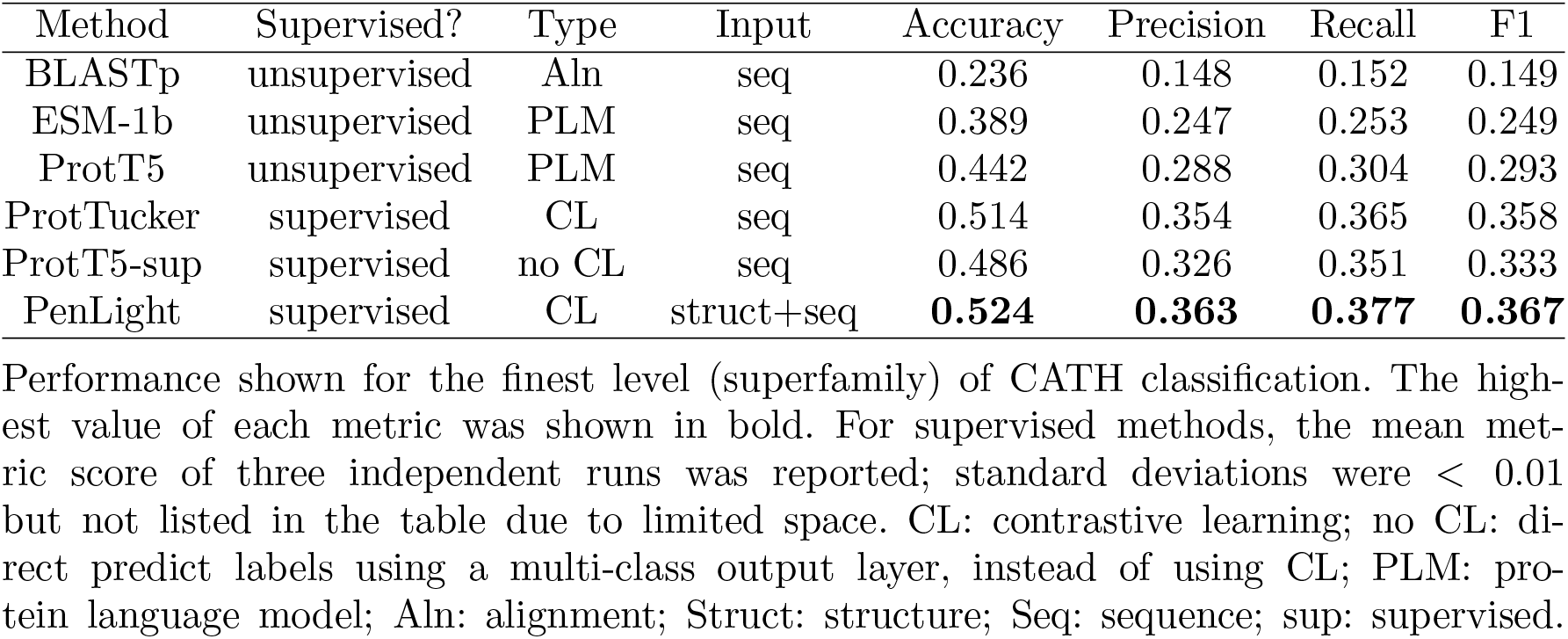
Performance on CATH structure classification.

| Method | Supervised? | Type | Input | Accuracy | Precision | Recall | F1 |
| --- | --- | --- | --- | --- | --- | --- | --- |
| BLASTp | unsupervised | Aln | seq | 0.236 | 0.148 | 0.152 | 0.149 |
| ESM-1b | unsupervised | PLM | seq | 0.389 | 0.247 | 0.253 | 0.249 |
| ProtT5 | unsupervised | PLM | seq | 0.442 | 0.288 | 0.304 | 0.293 |
| ProtTucker | supervised | CL | seq | 0.514 | 0.354 | 0.365 | 0.358 |
| ProtT5-sup | supervised | no CL | seq | 0.486 | 0.326 | 0.351 | 0.333 |
| PenLight | supervised | CL | struct+seq | **0.524** | **0.363** | **0.377** | **0.367** |

Performance shown for the finest level (superfamily) of CATH classification. The highest value of each metric was shown in bold. For supervised methods, the mean metric score of three independent runs was reported; standard deviations were $< 0.01$ but not listed in the table due to limited space. CL: contrastive learning; no CL: direct predict labels using a multi-class output layer, instead of using CL; PLM: protein language model; Aln: alignment; Struct: structure; Seq: sequence; sup: supervised.

### 3.2. Performance on Functional Annotations

After benchmarking PenLight on structure classification, we proceeded to evaluate PenLight’s ability to predict functional annotations. We used the structure dataset collected in Gligorijevic et al.,^19^ which contains 10,245 chains from the PDB database that have EC number annotations. The most specific (4th) level of EC numbers was used as the functional annotations to train and evaluate the models. The median number of samples per EC number is 12. The dataset was split into train, validation, and test sets with an approximate ratio 8:1:1, and the test set has no sequence sharing > 40% sequence identity to the training sequences.

Similar to the results of CATH classification, we also found that PenLight has learned embeddings that are discriminative between EC numbers (Fig. 2b). We compared PenLight with four state-of-the-art deep learning methods and found that PenLight achieved substantially higher performance (Table 2). PenLight first outperformed ProteInfer,^26^ DeepEC,^11^ and ProtTucker, three models that only take the amino acid sequence as input. PenLight also outperformed DeepFRI^19^ by a large margin, which is a GNN model that considers both the sequence and structure of the input protein but was trained using a supervised multi-class scheme. An ablation evaluation of PenLight showed that contrastive learning has led to better performance than the multi-class classification paradigm (PenLight(-) in Table 2).

**Table 2.** Performance on EC number prediction.

| Method | Type | Input | Cov | Accuracy | Precision | Recall | F1 | F1@Called |
| --- | --- | --- | --- | --- | --- | --- | --- | --- |
| DeepEC | CNN | seq | 0.34 | 0.287 | 0.466 | 0.326 | 0.361 | 0.737 |
| DeepFRI | GNN | str+seq | 0.60 | 0.442 | 0.451 | 0.353 | 0.380 | 0.432 |
| ProteInfer | CNN | seq | 0.42 | 0.367 | 0.538 | 0.414 | 0.448 | **0.758** |
| ProtTucker | CL | seq | 1.00 | 0.768 | 0.709 | 0.719 | 0.695 | 0.695 |
| PenLight(-) | no CL | str+seq | 1.00 | 0.676 | 0.604 | 0.609 | 0.585 | 0.585 |
| PenLight | CL | str+seq | 1.00 | **0.777** | **0.720** | **0.736** | **0.711** | 0.711 |

Performance shown for level 4 (most specific level) of EC number. The highest value of each metric was shown in bold. Coverage (Cov) is the fraction of test proteins for which a method makes a prediction. Proteins for which a method did not make a prediction (not called) will be counted as an incorrect prediction for metrics accuracy, precision, recall, and F1, but not for the F1@Called metric, which was calculated on called proteins of a method. The mean metric score of three independent runs was reported. Standard deviations were $< 0.01$ but not listed in the table due to limited space. CL: contrastive learning; Str: structure; Seq: sequence. no CL: direct predict labels using a multi-class output layer, instead of using CL.

Notably, ProteInfer, DeepEC, and DeepFRI all have a coverage (defined as the fraction of test proteins for which the method made predictions) lower than PenLight because they only predict EC numbers for a query protein when the predicted score passes a predefined confidence threshold (we say it is a “called protein” hereafter). In contrast, PenLight always predicts for the query protein by transferring the known EC numbers from the top-1 closet lookup protein, thus having a prediction coverage of 1.0. To make a fair comparison, we also restricted the evaluation on called proteins for those baselines, i.e., not counting non-called proteins as wrong predictions. We found that in this case PenLight still had a higher F1 score than DeepFRI (‘F1@Called’ column in Table 2). DeepEC and ProteInfer achieved a slightly higher F1 than PenLight but at an expense of much lower (< 0.5) coverage. Despite PenLight always predicting for every protein by transferring from the top-1 closest lookup protein, it is also possible to introduce a confidence threshold for PenLight, similar to those in our baseline methods, which will be demonstrated in the next section. Overall, the performance improvements achieved by PenLight in this task again demonstrated the advantages of integrating structure data and contrastive learning for protein function prediction.

### 3.3. Analyses of PenLight’s high coverage and high accurate predictions

Here, we further dissect the relationship between PenLight’s prediction accuracy and coverage. We first performed a detailed stratified comparison of prediction accuracy on the EC number prediction task. Specifically, we plotted the proportion of correct, incorrect and not called predictions of PenLight, ProteInfer, and DeepFRI at each EC number level (Figures 3a-c). ProteInfer had quite stable prediction accuracies ~0.4) across the four levels but failed to predict the EC numbers for approximately 57% of proteins. For DeepFRI, as the EC number levels become more specific (from level 1 to 4), both its prediction accuracy and coverage dropped, likely due to proteins being more similar in sequence at higher EC number levels, and it is more challenging to distinguish their differences in function. In contrast, PenLight had an accuracy > 0.75 for all four levels while maintaining a 100% coverage. The major reason for the high accuracy and high coverage of PenLight is the contrastive learning and the lookup strategy for making predictions. Methods like ProteInfer formulated the CATH code or EC number classification as a supervised multi-class classification problem and predict the class probabilities for thousands of classes using the single final layer in the neural network. This strategy inevitably suffers from the class size imbalance in the training data, and the ambiguity in the output layer is easily scaled up with the number of classes (e.g., thousands of EC numbers or CATH codes). On the contrary, PenLight first applied contrastive learning to learn discriminative embeddings with respect to the functional or structural annotations, reducing the ambiguity between positive and negative data points (Fig. 2). PenLight then enumerated all proteins in the lookup set and identified the protein with the closest distance to the query protein. This similarity search process treats the distance to every lookup protein equally, without down-weighting any under-represented classes. Therefore, PenLight was able to accurately predict the labels even for under-represented EC numbers, where supervised-learning approaches often have large uncertainties. In our tests, we observed that when predicting for EC numbers that have only < 10 proteins in the training set, PenLight achieved an accuracy of 0.8 while ProteInfer, DeepEC, and DeepFRI only had an accuracy of ~ 0.6.

**Fig. 3.**
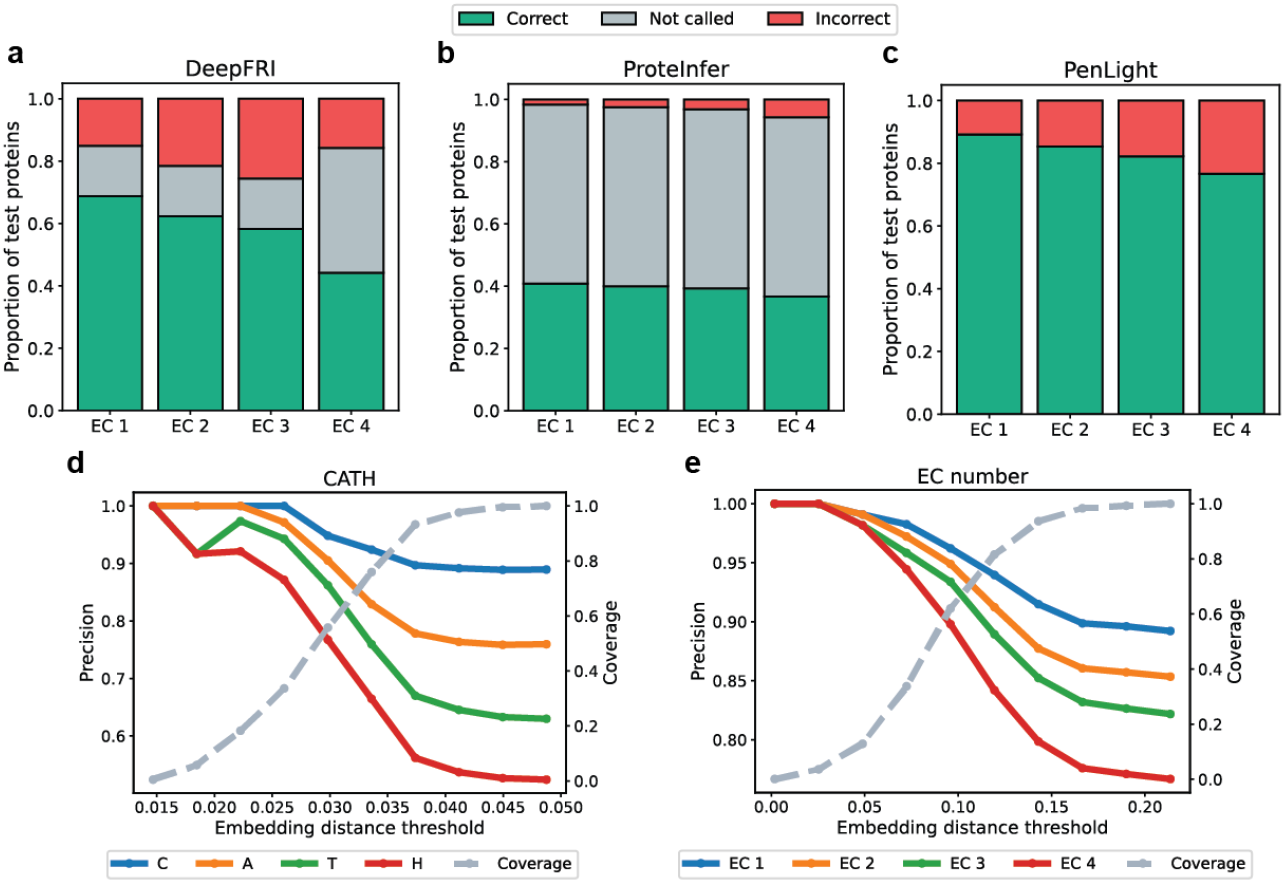
PenLight achieved prediction coverage and accuracy. (a-c) Stacked bar plots of DeepFRI, ProteInfer and PenLight that visualized the fractions of correct, incorrect, and not called predictions at the four levels of EC numbers. (d-e) PenLight’s prediction coverage and precision as a function of the embedding distance threshold. Here PenLight predicts the CATH code (d) or EC number (e) for a protein if its closest embedding distance to the lookup set protein is below a given threshold (called a hit). Coverage is defined as the fraction of hit in all test proteins, and precision is defined as the fraction of correct predictions for the hit proteins.

We next explore the possibility of introducing a confidence threshold into PenLight, similar to the E-value cutoff used in BLAST. A natural choice is to impose a cutoff on the Euclidean embedding distance, i.e., making predictions only when the query protein’s closest distance to lookup proteins is below the cutoff. We thus varied the distance cutoff and evaluated how the prediction precision and coverage would change as the cutoff was changing. As expected, we observed that PenLight had a high prediction precision for the CATH task when the cutoff was very stringent (smaller values) since the model was confident in this regime of cutoff values (Fig. 3d). On the other hand, when the cutoff became more tolerant (larger values), the precision started to drop but the prediction coverage gradually increase. Similar trends were observed for EC task as well (Fig. 3e). Overall, this analysis validated that PenLight’s embedding distance was correlated with prediction accuracy, and a cutoff can be used to tradeoff the prediction precision and coverage, depending on the practical use case (e.g., accurate annotations or data explorations).

Finally, we performed a t-SNE visualization to see whether PenLight has learned meaningful representations in terms of structural and functional similarity. We observed that, on the CATH task, the embedding space learned by PenLight was a more consistent with the CATH hierarchy, where the ProtT5’s embeddings did not capture the structural similarities of CATH classes (Figs. 4a) while PenLight’s embeddings showed separated grouping structures consistent with the first level of CATH classification (Fig. 4b). Similarly, on the EC number task, we found that PenLight’s embedding space showed clustering patterns more consistent with six major enzyme groups than the ESM-1b model (Figs. 4c-d).

**Fig. 4.**
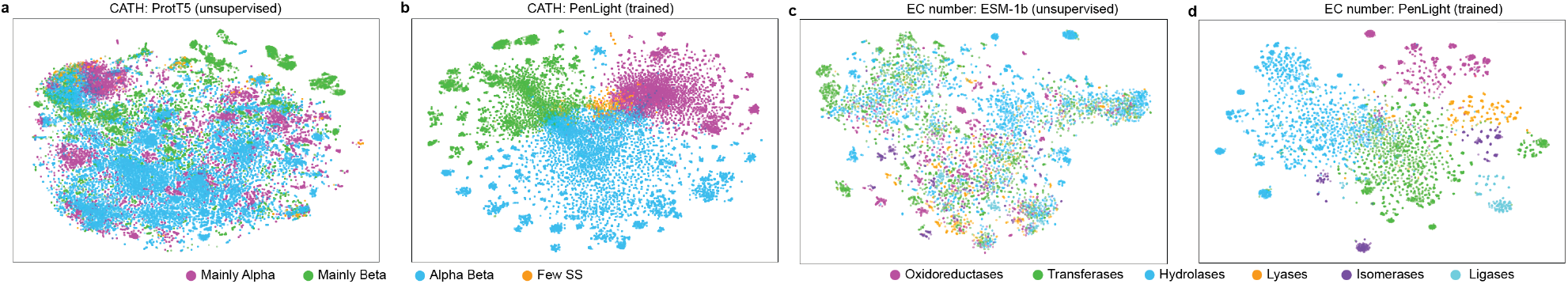
t-SNE visualizations. Embedding space learned by PLMs and PenLight on (a) the CATH dataset and (b) the EC number dataset. Two PLMs (ProtT5 for CATH and ESM-1b for EC) were shown for comparison. One point represents a protein. Points were colored according to their assigned label at the first level of CATH class or EC number.

## 4. Conclusions

We described PenLight, a general deep learning framework that predicts protein structural and functional annotations. PenLight integrates 3D protein structures and protein language model embeddings with a structure-aware graph neural network (GNN). To learn protein representations that capture meaningful structural or functional similarities, PenLight used a contrastive learning strategy to train the GNN. We showcase PenLight’s applicability using both structural and functional annotation tasks, and the experiment results suggested that PenLight outperformed several state-of-the-art methods in predicting the CATH structure hierarchy and enzyme class of proteins. As a general framework, PenLight can be extended to other protein annotation tasks as well, such as gene ontology classification. Recent progress in the graph deep learning community, including equivariant graph neural network,^27^ can also be integrated with PenLight to enable better structure-based protein annotation.

## Acknowledgements

This work was supported by the 2022 Seed Grant Program of the Molecule Maker Lab Institute, an NSF AI Institute (grant no. 2019897).

